# Kinetic profiling of metabolic specialists demonstrates stability and consistency of in vivo enzyme turnover numbers

**DOI:** 10.1101/767996

**Authors:** David Heckmann, Anaamika Campeau, Colton J. Lloyd, Patrick Phaneuf, Ying Hefner, Marvic Carrillo-Terrazas, Adam M. Feist, David J. Gonzalez, Bernhard O. Palsson

## Abstract

Enzyme turnover numbers (*k*_cat_s) are essential for a quantitative understanding of cells. Because *k*_cat_s are traditionally measured in low-throughput assays, they are often noisy, non-physiological, inconsistent, and labor-intensive to obtain.

We use a data-driven approach to estimate in vivo *k*_cat_s using metabolic specialist *E. coli* strains that resulted from gene knockouts in central metabolism followed by metabolic optimization via laboratory evolution. By combining absolute proteomics with fluxomics data, we find that *in vivo k*_cat_s are robust against genetic perturbations, suggesting that metabolic adaptation to gene loss is mostly achieved through other mechanisms, like gene-regulatory changes. Combining machine learning and genome-scale metabolic models, we show that the obtained *in vivo k*_cat_s predict unseen proteomics data with much higher precision than *in vitro k*_cat_s. The results demonstrate that *in vivo k*_cat_s can solve the problem of noisy and inconsistent parameterizations of cellular models.

## Introduction

Enzyme catalytic rates are crucial for understanding many properties of living systems like growth, proteome allocation, stress, and dynamic responses to perturbation. The turnover number of an enzyme, *k*_cat_, describes the maximal rate at which an enzyme’s catalytic site can catalyze a reaction. Knowledge of *k*_cat_ has traditionally been a bottleneck in the quantitative understanding of cells, mainly because *k*_cat_s have historically been obtained in labor-intensive, low-throughput in vitro assays. The substantial effort required for in vitro assays is likely the reason why, even in model organisms, only a small fraction of cellular enzymes has a measured *k*_cat_^1^. Furthermore, in vitro *k*_cat_ estimates are frequently very inconsistent when different literature sources are compared^2^, probably because in vitro conditions do not mimic in vivo conditions, are affected by post-translational modifications, and can be biased by experimental batch effects.

In order to address the problems of low-throughput acquisition and in vivo-in vitro discrepancies, Davidi et al.^3^ combined proteomics data and flux predictions to estimate in vivo turnover numbers based on apparent catalytic rate (*k*_app_). Davidi et al.^3^ integrated published *E. coli* proteomics data sets with in silico flux predictions in multiple growth conditions and showed that the resulting maximum apparent catalytic rate (*k*_app,max_) across growth conditions is significantly correlated with in vitro *k*_cat_s. Thus, *k*_app,max_ has the potential to overcome the problem of inconsistencies, in vitro-in vivo discrepancies, and batch effects from which in vitro *k*_cat_s suffer. However, it is unclear if *k*_app,max_ is a stable system parameter that is robust to perturbation, and how much experimental procedures bias the estimation of *k*_app,max_: absolute proteomic quantification techniques are still suffering from high variation and previous estimates of *k*_cat_ were based on in silico flux predictions rather than ^13^C fluxomics data. Furthermore, *k*_cat_ is expected to scale with growth rate^4^. As many experimental conditions in the literature data used in Davidi et al.^3^ resulted in low growth rates, the effective number of data sets contributing to *k*_app,max_ is low. Finally, if *k*_app,max_ is a useful estimator of in vivo *k*_cat_, it should improve the predictive capability of metabolic models on data that was not used to obtain *k*_app,max_, i.e., the performance on a test set.

Here, we present a new approach for estimating *k*_cat_ in vivo (Figure 1). We combined proteomic profiling with fluxomics data to estimate in vivo *k*_cat_s in *E. coli* strains that have undergone strong physiological perturbations via knockout of metabolic genes. To obtain strains with high growth rates for which *k*_app_ approaches *k*_app,max_, adaptive laboratory evolution^5^ was used on the metabolic knockout strains. We profiled 21 strains, representing metabolic specialists with diverse flux profiles that are able to obtain high growth rates^6–9^. With this data-driven approach, we show that in vivo *k*_cat_s are stable and robust to genetic perturbations, and that they can be used in genome-scale models to obtain a high predictive performance for unseen protein abundance data.

**Figure 1:**
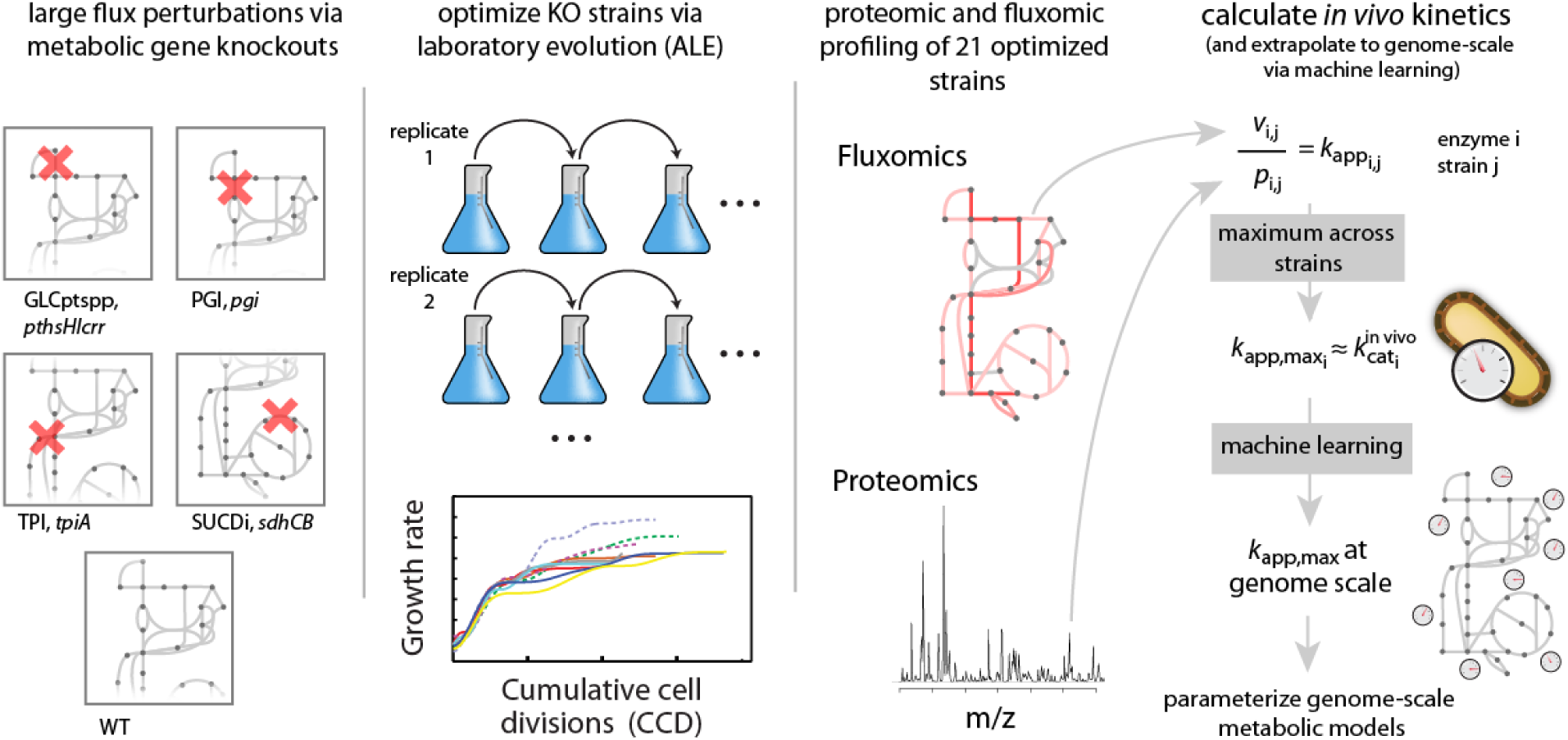
Approach for obtaining *k*_cat_ in vivo from metabolic specialists: Knock out of enzymes in central metabolism was followed by adaptive laboratory evolution (ALE) to obtain 21 strains that had diverse flux profiles, while achieving high growth rates^6–9^. Fluxomics and proteomics data was then integrated for the evolved strains to obtain the maximum *k*_app_ across the 21 strains (*k*_app,max_) for each enzyme that could be mapped uniquely. The obtained *k*_app,max_ vector was then extrapolated to genome scale via supervised machine learning and used to parameterize genome-scale metabolic models. The resulting genome-scale models were then validated on unseen proteomics data.

## Results

### Quantifying in vivo kinetics in metabolic specialists

In theory, *k*_app,max_ will approach *k*_cat_ in vivo if a condition is found in which the respective enzyme is utilized at full efficiency. In order to achieve strong genetic perturbations of enzyme usage, we used gene knockout (KO) strains for the PTS system (*ptsHIcrr*^*6*^), the phosphoglucose isomerase (*pgi*^*8*^), triosephosphate isomerase (*tpiA*^*7*^), and succinate dehydrogenase (sdhCB^9^). As *k*_app_ increases with growth rate^4^, we used KO strains that were optimized for growth on glucose minimal medium via adaptive laboratory evolution (ALE)^6–9^ experiments. In addition to these KO strains, we utilized a wild type MG1655 strain that was subjected to ALE^6–10^. As evolution is not a deterministic process, ALE endpoints differ in genotype, and we included a total of 21 strains that resulted from replicates of ALE experiments (i.e., four endpoint strains for *ptsHIcrr*, eight for *pgi*, four for *tpiA*, three for *sdhCB*, and two WT controls) and that were representative for the respective endpoint population. We subjected the selected strains to genome sequencing and used the resulting sequences as reference proteomes in LC-MS/MS proteomics (see Methods). Absolute quantification of biological duplicates was achieved via calibration to the UPS2 standard and the top3 metric^11,12^, which estimates protein abundance based on the average intensity of the three best ionizing peptides. Measured protein abundances show a median *R*^2^ of 0.91 between biological replicates, and a median number of 2076 proteins were detected per strain (Supplementary Table 1). The obtained protein abundance vectors cluster by the genetic background of the strain used for ALE (Supplementary Figure 1), indicating that protein levels have adjusted in a specific pattern to compensate for the respective gene KO (see ^6–9^ for details on the transcript level).

### Gene KO and ALE cause diversity in enzyme usage

We integrated the measured protein abundances with ^13^C MFA fluxomics data^6–9^ to calculate apparent catalytic rates in the 21 strains as the ratio of flux and protein abundance. Like in Davidi et al.^3^, we only calculated *k*_app_ for homomeric enzymes and reactions that are not catalyzed by multiple isoenzymes to allow a specific mapping of proteins to reactions. This approach resulted in a median number of 258 enzymes per strain for which we were able to calculate *k*_app_. The resulting apparent catalytic rates largely cluster by the genotype of the KO strain (Figure 2A), confirming that enzyme usage was indeed perturbed by the respective KOs. Across the 21 strains, the maximum observed *k*_app_ of an enzyme is on average 4.4 times larger than the smallest observed *k*_app_ (Figure 2B), indicating that considerable variation in enzyme usage was caused by the metabolic gene knockout. To exclude the possibility that experimental variation causes this apparent diversity in enzyme usage, we compared the standard deviation of *k*_app_ in biological replicates (mean on log_10_ scale = 0.07) to the standard deviation measured across the 21 strains (mean on log_10_ scale = 0.18). We found that the variation caused by KOs and ALE is significantly larger than that caused by experimental variation (p<2e-16, Wilcoxon rank sum test).

**Figure 2:**
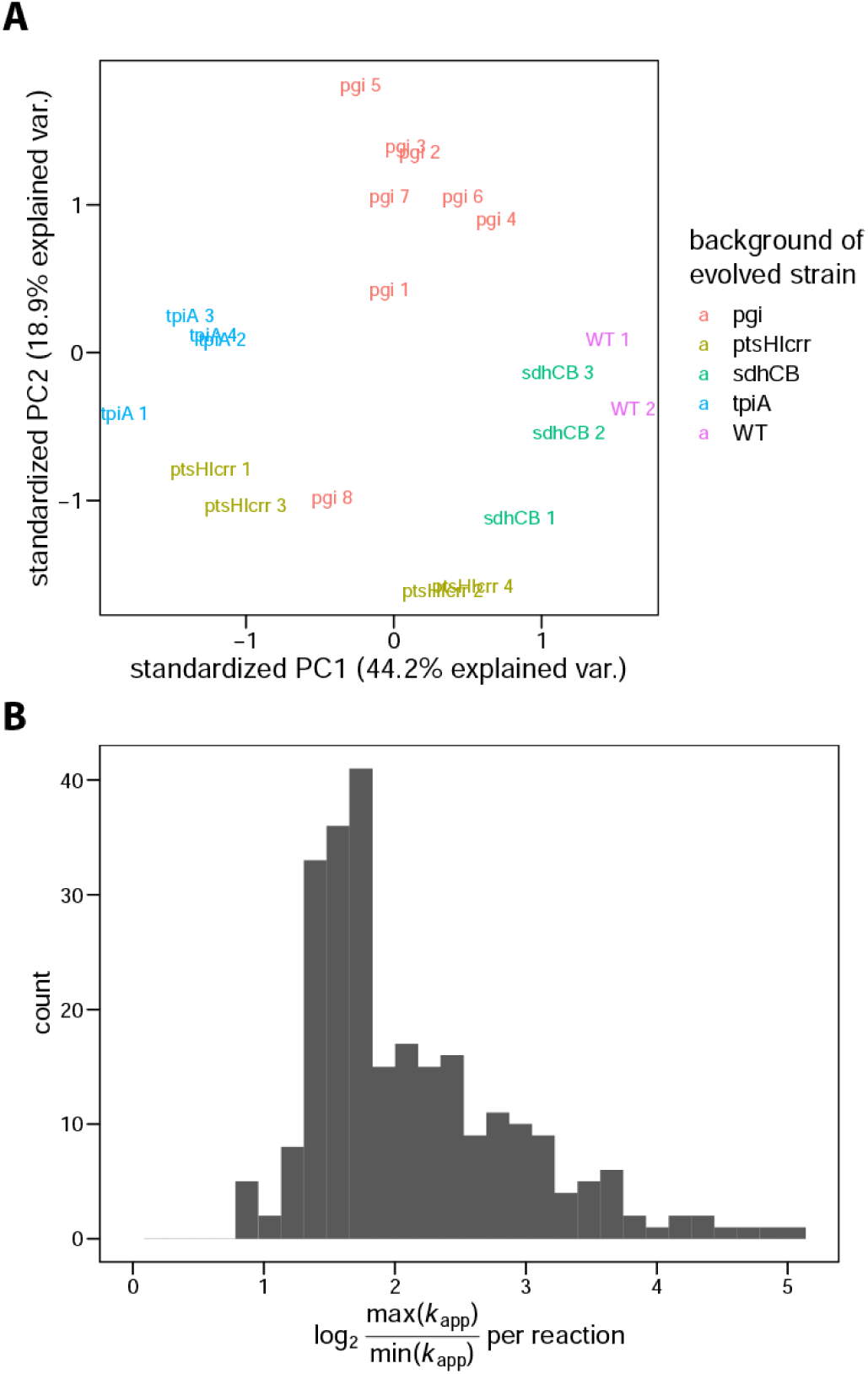
Apparent catalytic rates cluster by genetic background and exhibit diversity across strains. (A) Data on *k*_app_ in each of the 21 strains projected onto the first two principal components. Only reaction-strain combinations for which *k*_app_ was available in all strains were used, resulting in 214 reactions used in the analysis. Data was centered and scaled before conducting principal component analysis. (B) Distribution of ranges of *k*_app_ across reactions. The log_2_ of the ratio between the highest and the lowest *k*_app_ per reaction is shown.

### In vivo turnover numbers are stable and consistent

We estimated in vivo *k*_cat_ for a given enzyme as the maximum of *k*_app_ (*k*_app,max_) in the 21 KO strains. This was similar to Davidi et al.^3^, who estimated in vivo *k*_cat_s as the maximum *k*_app_ over different growth conditions. Due to incomplete substrate saturation and backward flux, the apparent catalytic rate of an enzyme is smaller than the in vivo *k*_cat_. It is thus unclear if *k*_app,max_ is a stable property of the system that can be used in metabolic models to give reliable predictions. Furthermore, absolute proteomics data and fluxomics data come with significant experimental uncertainties and biases that could prevent *k*_app,max_ from being useful in modeling applications.

Even though our protocol perturbed enzyme *k*_app_ via gene KO and ALE, whereas Davidi et al.^3^ used differences in growth conditions to achieve variation in enzyme usage, we found a very high agreement between *k*_app,max_ from the two sources (*R*^2^ = 0.9, Figure 3A). We used a parametric bootstrap procedure to quantify the uncertainty in our *k*_app,max_ estimations (see Methods). We found that 42% (88 out of 210) of comparable values estimated by Davidi et al.^3^ fall into the 95% confidence intervals of the *k*_app,max_ values obtained in this study. A clear outlier is the reaction FAD reductase (FADRx, Figure 3A). This discrepancy is caused by the different methods of flux estimation: while protein abundances of FAD reductase are relatively similar in the respective conditions for which the maximum *k*_app_ was measured (protein abundance is two times lower in Davidi et al.^3^), flux through the FADRx reaction in parsimonious FBA^13^ is one thousand times higher than the flux estimated in ^13^C MFA.

**Figure 3:**
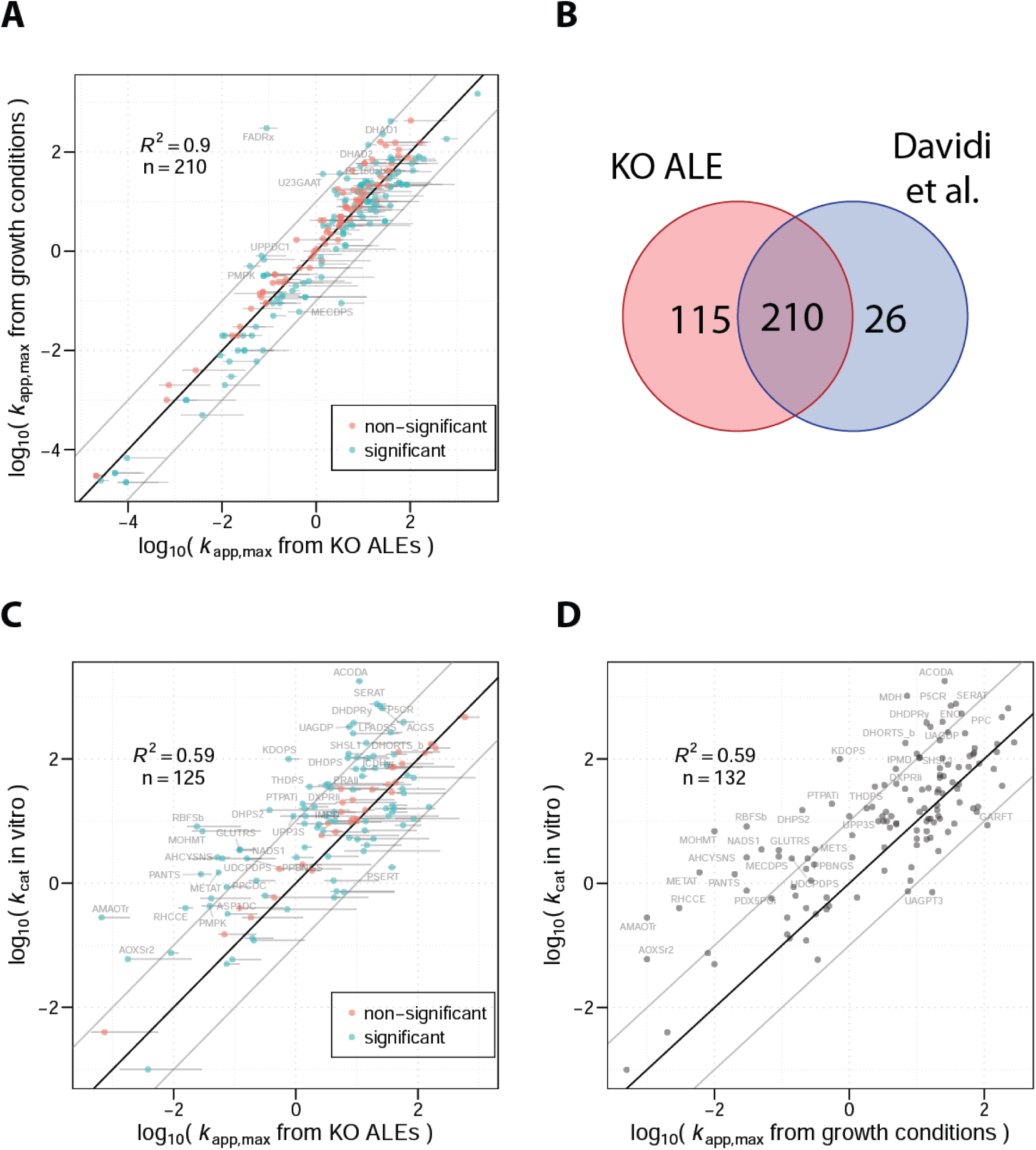
Estimates of in vivo turnover numbers are consistent. (A) Comparison between *k*_app,max_ obtained from KO strains (this study) and *k*_app,max_ from growth conditions (Davidi et al.^3^). (B) Number of reactions for which *k*_app,max_ was obtained in KO strains (this study) and varying growth conditions (Davidi et al.^3^). (C) Comparison between *k*_app,max_ obtained from KO strains and in vitro *k*_cat_s. (D) Comparison between *k*_app,max_ obtained from KO strains and in vitro *k*_cat_s^3^. Horizontal lines are 95% confidence intervals determined by 500 parametric bootstrap samples (see Methods). Points are marked red when the compared value falls into the 95% confidence interval of *k*_app,max_ and are labelled with reaction IDs as given in the *i*JO1366 reconstruction^15^ if the values differ by more than one order of magnitude. Data on *k*_cat_ in vitro shown in panel (C) and (D) was taken from Davidi et al.^3^ to allow for comparison between the studies.

It is worth noting that the mutations observed in the ‘ALE strains’ are mostly regulatory in nature, with almost no structural changes in the homomeric enzymes examined in this study (see Supplementary Data 1 and ^6–9,14^ for details). One exception is the enzyme isocitrate dehydrogenase, which has shown a very high level of convergence for a coding sequence mutation (R395C) in seven out of the eight evolved *pgi* KO strains. We found no significant difference in *k*_app,max_ compared to the *k*_app,max_ of isocitrate dehydrogenase reported by Davidi et al.^3^ (p=0.28, parametric bootstrap), suggesting that the structural mutation does not increase the in vivo catalytic efficiency.

While *k*_app,max_ from KO strains is very consistent with *k*_app,max_ from different growth conditions, the correlation with *k*_cat_ in vitro is significantly lower (*R*^2^ = 0.59, Figure 3C), and only 26% (32 out of 125) of the in vitro values fall into the 95% confidence intervals of *k*_app,max_. A similar low correlation with in vitro *k*_cat_ was found in the *k*_app,max_ estimates published by Davidi et al.^3^ (*R*^2^ = 0.59, Figure 4D).

**Figure 4:**
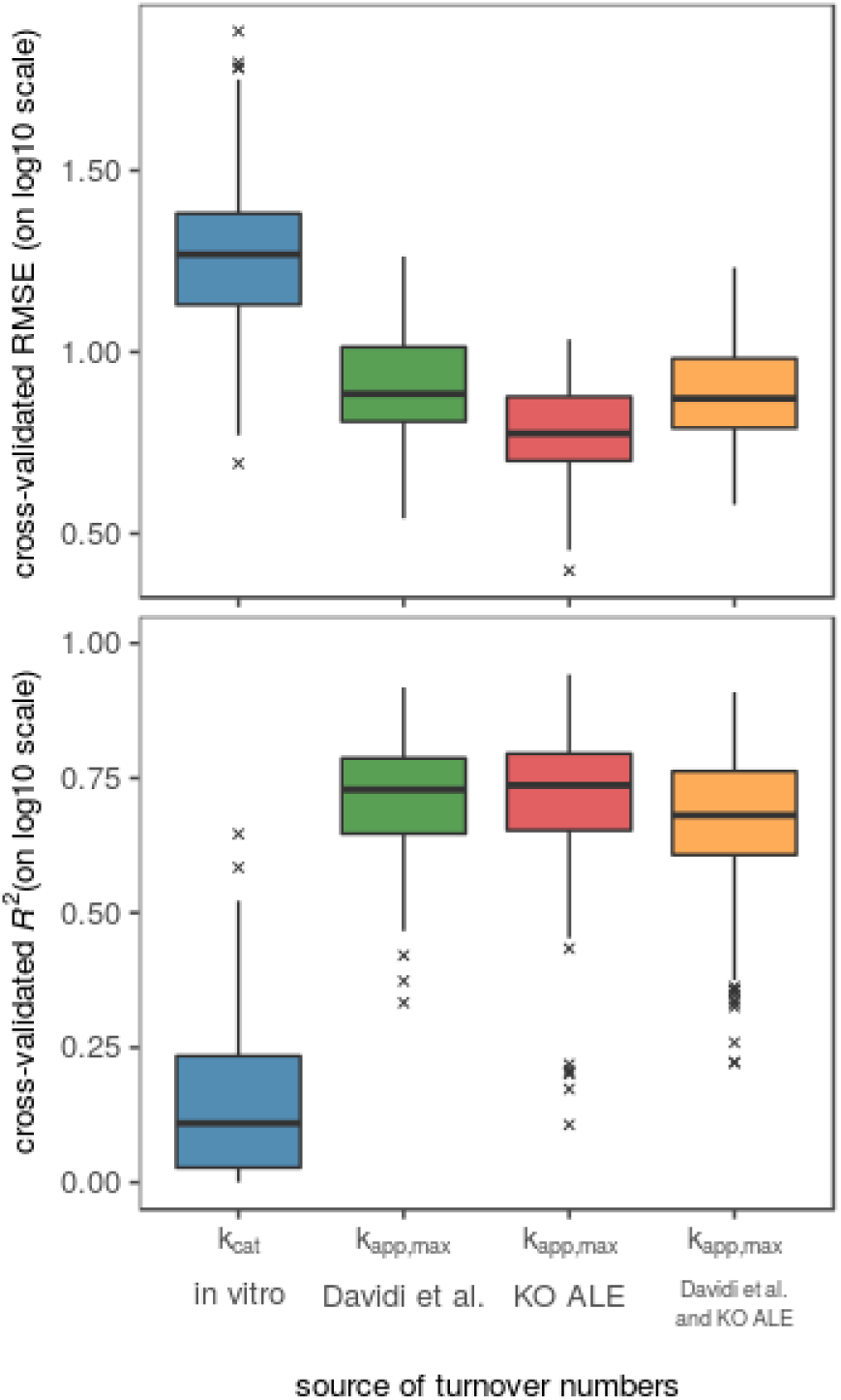
Performance of machine learning models on different sources of turnover numbers. The performance is estimated in five-times repeated five-fold cross-validation in elastic net, random forest, and a neural network^16^ (see Methods). Data for *k*_cat_ in vitro was taken from Heckmann et al.^16^.

In summary, although we obtained *k*_app,max_ from a genetic perturbation rather than variation in growth conditions, used ^13^C fluxomics data instead of in silico flux, and despite proteomics and flux data being subject to significant noise, we found very high agreement between *k*_app,max_ from the two sources.

### Using machine learning to extrapolate to the genome scale

The problem of low coverage that is associated with *k*_cat_ in vitro is also present in *k*_app,max_: not all protein abundance can be mapped to enzymes uniquely, and proteomics experiments still suffer from coverage issues. The final set of *k*_app,max_ values includes 325 enzymes (Figure 3B). This coverage is 27% higher than that found in Davidi et al.^3^, mostly because we used ^13^C fluxomics data that tends to have a higher sensitivity than the in silico method (parsimonious FBA^13^) used by Davidi et al.^3^. In order to validate the estimated in vivo turnover numbers in a genome-scale model that contains over three thousand direction-specific reactions, we first needed to extrapolate the data to the genome scale. We used supervised machine learning on a diverse enzyme data set^16^ that includes data on enzyme network context, enzyme 3D structure, and enzyme biochemistry to achieve this goal. An ensemble model of an elastic net, random forest, and neural network^16^ showed good performance in cross-validation for the in vivo turnover numbers, where the highest performance was achieved for *k*_app,max_ that was obtained from the 21 KO strains (Figure 4). Taking the maximum of *k*_app,max_ from this study and that of Davidi et al.^3^ did not improve model performance, even though it resulted in the largest training set.

### Validation of turnover numbers in mechanistic models

The enzyme turnover number is a major determinant of gene expression levels as it sets a lower limit on the enzyme concentration required to maintain a given flux. Turnover numbers are successfully used in genome-scale metabolic models to constrain metabolic fluxes by a limited cellular protein budget^17–19^ or the balance of translation and dilution of proteins^20–22^. The *k*_app,max_ obtained from diverse growth conditions was previously used successfully in genome-scale metabolic models, showing that the performance of protein abundance predictions of models using *k*_app,max_ was significantly higher than that of models using in vitro *k*_cat_s^16^. A major drawback of this analysis lies in the validation of the metabolic model which used data^23^ that was also utilized in obtaining *k*_app,max_^3^, posing the risk of circular reasoning through data leakage. If *k*_app,max_ is a stable property of in vivo enzyme catalysis, it is expected to yield a high performance in metabolic models on unseen data, i.e., *k*_app,max_-based models should generalize well.

To test this hypothesis, we parameterized two genome-scale modeling algorithms of proteome-limited metabolism, a MOMENT^17^ and an ME model^20^, with *k*_app,max_ obtained from KO strains. We then used the model to predict enzyme abundance data under various growth conditions published by Schmidt et al.^23^, a data set that was not used to obtain *k*_app,max_ in this study. For comparison, we included model parameterization based on *k*_cat_ in vitro, with *k*_app,max_ from Davidi et al.^3^, and the maximum of *k*_app,max_ obtained in this study and that of Davidi et al.^3^. We found that the performance of *k*_app,max_ from KO strains on the Schmidt et al.^23^ data is very similar to that of Davidi et al.^3^: the average median root mean squared error (RMSE) on log_10_ scale is 4% higher for the MOMENT model and 12% lower for the ME model, even though the Schmidt et al. data^23^ was used to obtain *k*_app,max_ in Davidi et al.^3^ (Figure 5, Supplementary Figure 2). This good performance on unseen data confirms that in vivo *k*_cat_ are stable against genetic perturbation and consistent across experimental protocols.

**Figure 5:**
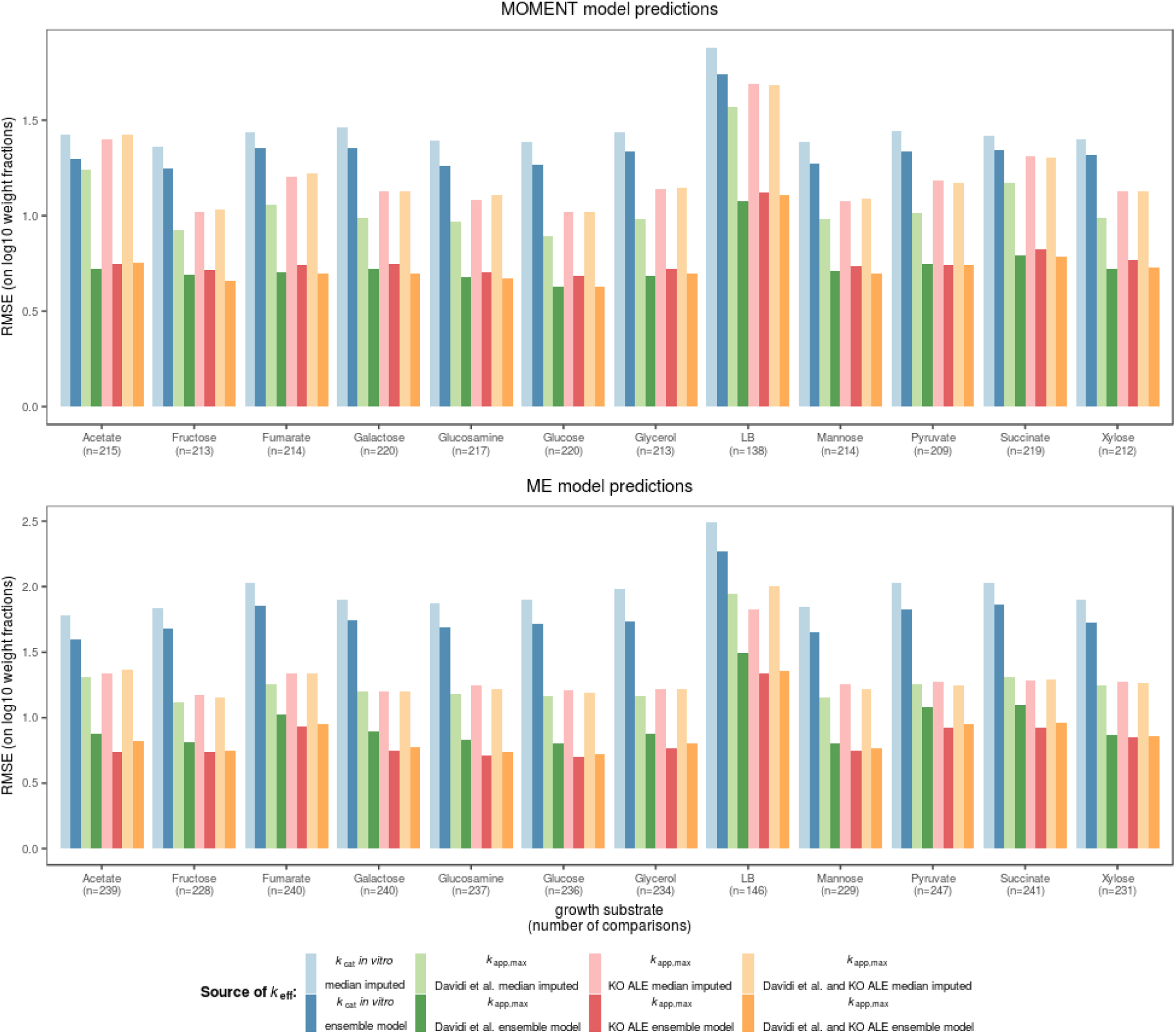
Performance in mechanistic models of proteome allocation. The MOMENT algorithm and a ME model were parameterized with different sources of turnover numbers. Growth on different carbon sources was simulated with the two algorithms and the predicted relative protein weight fractions of metabolic enzymes were compared to proteomics data in the respective growth condition^23^ using the root mean squared error (RMSE) on log_10_ scale.

We further found that *k*_app,max_ outperforms *k*_cat_ in vitro in MOMENT and ME models across all growth conditions (Figure 5). When comparing median-imputed *k*_cat_ parameterizations to those using supervised machine learning, we found that machine learning reduces RMSE on log_10_ scale by 38% for *k*_app,max_ and 10% for *k*_cat_ in vitro, confirming the utility of this approach^16^.

## Discussion

A large-scale characterization of the kinetic parameters that govern metabolism, termed the kinetome^1^, has been a major hurdle in our quantitative understanding of cellular behavior^1,24,25^. Previous efforts to use *k*_cat_, which represents a major fraction of the kinetome, at the genome scale either utilized in vitro data^17,19^ or fitted kinetic parameters to physiological data^4,26,27^. While in vitro *k*_cat_s suffer from inconsistencies, low-throughput, high noise, and missing in vivo effects, parameter fitting is frequently under-determined and leads to non-unique solutions that can not be expected to generalize well when used in new conditions. The use of proteomics data and flux predictions on homomeric enzymes^3^, for which proteome abundances can be assigned uniquely, is a promising approach that could solve many shortcomings of in vitro data and fitting approaches. While it was shown that this approximation of in vivo *k*_cat_, *k*_app,max_, exhibits a decent correlation with *k*_cat_ in vitro, it is unclear if *k*_app,max_ captures an upper bound on enzymatic rate that is stable with respect to genetic perturbations and consistent across experimental procedures. These properties are prerequisites for the application of *k*_app,max_ in metabolic models.

We found that in vivo turnover numbers that are obtained from KO strains are surprisingly consistent between very different protocols (Figure 3). Specifically, the protocol we used to obtain *k*_app,max_ shows the following differences compared to that of Davidi et al.^3^: (1) *k*_app_ is not perturbed by growth conditions, but by genetic KOs; (2) we used ^13^C MFA fluxomics data instead of in silico data from parsimonious FBA; (3) we utilized proteomics data that was obtained with a single LC-MS/MS protocol, avoiding batch effects; and (4) all data was obtained under batch growth that promotes high growth rates, increasing *k*_app_^4^. Given these differences in the two approaches to obtain *k*_app,max_, the high agreement between the two methods indicates a high stability and consistency of in vivo *k*_cat_s.

The high stability of in vivo *k*_cat_s indicates that the adaptation of the strains during ALE does not lead to drastic increases in in vivo *k*_cat_s. This hypothesis is supported by the relatively low number of convergent mutations in the coding regions of enzymes (Supplementary Data 1). Short term metabolic evolution appears to be governed by changes in gene regulation, rather than changes in enzyme efficiencies, at least in the case of the homomeric enzymes investigated in this study.

Why does *k*_app,max_ exhibit a high consistency, where, in contrast, in vitro *k*_cat_s often show a low agreement between different sources^2^? One reason might lie in the avoidance of batch effects: in vitro *k*_cat_s are typically obtained individually in enzyme-specific assays, whereas *k*_app,max_ used a small number of proteomics and flux data that were ideally obtained on the same instruments, thus avoiding batch effects. Furthermore, there is some indication that metabolite levels in vivo tend to saturate many enzymes^28^, allowing for conditions of high enzyme saturation to be found even with a relatively small number of system perturbations.

Some sources of uncertainty remain in the *k*_app,max_ values presented in this study. The ^13^C MFA data that we used was obtained for the endpoint populations of the respective ALE experiments^6–9^, whereas we used clonal samples for proteomics experiments. While we chose clones that represented the most dominant mutations found in the endpoint populations, flux distributions could be affected by uncommon mutations. Furthermore, ^13^C MFA data can yield high coverage^29^, but it still relies heavily on the quality of the underlying network model, which could bias analyses.

Because not all enzymes can be mapped to reaction uniquely and proteomics data still suffers from incomplete coverage, *k*_app,max_ has a low coverage of the metabolic network and can not be readily used in genome-scale models. Based on mechanistic knowledge of factors that shape enzyme turnover numbers^2,30,31^, supervised machine learning was previously used successfully to extrapolate in vivo *k*_cat_s to the genome scale^16^. We find a slightly lower error in cross validation on *k*_app,max_ obtained from KO strains compared to *k*_app,max_ from varying growth conditions^3^; this slight increase in performance may lie in the increased size of the training set, as we were able to obtain 38% more *k*_app,max_ values due to the use of ^13^C MFA data. This finding is consistent with previously computed learning curves of *k*_app,max_ on the Davidi et al. data set^3^ that showed that a domain of diminishing returns in model performance is reached with respect to the size of the training set^16^.

We find that metabolic models that are parameterized with the *k*_app,max_ values we obtained from KO strains lead to very good predictive performance on unseen proteomics data. This performance in mechanistic models supports the hypothesis that *k*_app,max_ indeed represents a stable property of the system, i.e. *k*_cat_ in vivo. Thus, *k*_app,max_ can enable genome-scale metabolic models that generalize well to unseen conditions.

While kinetic parameters remain difficult to obtain, the stable and consistent properties of in vivo *k*_cat_s support the notion that these parameters can improve the predictive capabilities of metabolic models significantly, and thus enable better quantitative understanding of the cell. Finally, the high stability of in vivo *k*_cat_s suggests that short-term metabolic evolution is governed by changes in gene expression, rather than adaptation at the level of enzyme kinetics.

## Methods

### Strain genomic sequencing

Genomic DNA of ALE endpoint clones was isolated using bead agitation in 96-well plates as outlined previously^32^. Paired-end whole genome DNA sequencing libraries were generated with a Kapa HyperPlus library prep kit (Kapa Biosystems) and run on an Illumina HiSeq 4000 platform with a HiSeq SBS kit, 150 bp reads. The generated DNA sequencing fastq files were processed with the breseq computational pipeline (version 0.32.0)^33^ and aligned to an *E. coli* K12 MG1655 reference genome^34^ to identify mutations. DNA-seq quality control was accomplished using the software AfterQC (version 0.9.7)^35^.

Clones were chosen in order to represent the high-frequency alleles found in the end-point populations of the respective ALE experiments. DNA sequences were used to create reference proteomes for proteomics experiments described below.

### Sample preparation

For each strain, 3 ml of culture was grown overnight at 37°C with shaking in M9 medium^36^ (4g Glucose l^-1^) with trace elements^37^, and then passed twice the following days in 15ml of media at 37°C from OD 0.05-0.1 to OD 1.0-1.5. For the experiment, 100 mL of culture with initial OD600 = 0.1 was grown in flasks with stirring in a water bath at 37°C. When cultures reached OD600 = 0.6, 40mL of culture was collected and immediately put on ice. The cells were pelleted by centrifuge at 5000 rpm at 4°C for 20 minutes. Cell pellets were then washed with 20 mL of cold PBS buffer three times and centrifuged at 5000 rpm for 20min at 4°C. Pellets were transferred into 1.5 mL micro centrifuge tubes and centrifuged at 8000rpm at 4°C for 10 minutes. Remaining PBS buffer was removed and pellets of proteomic samples were frozen at −80C.

### Sample lysis for proteomics

Frozen samples were immersed in a lysis buffer comprised by 75 mM NaCl (Sigma Aldrich), 3% sodium dodecyl sulfate (SDS) (Fisher Scientific), 1 mM sodium fluoride (VWR International, LLC), 1 mM β-glycerophosphate (Sigma Aldrich), 1 mM sodium orthovanadate, 10 mM sodium pyrophosphate (VWR International, LLC), 1 mM phenylmethylsulfonyl fluoride (Fisher Scientific), 50 mM HEPES (Fisher Scientific) pH 8.5, and 1X cOmplete EDTA-free protease inhibitor cocktail. Samples were subjected to rapid mixing and probe sonication using a Q500 QSonica sonicator (Qsonica) equipped with 1.6 mm microtip at amplitude 20%. Samples were subjected to three cycles of 10 seconds of sonication followed by 10 seconds of rest, with a total sonication time of 50 seconds.

### Protein Abundance Quantitation

Total protein abundance was determined using a BCA Protein Assay Kit (Pierce) as recommended by the manufacturer.

### Peptide Isolation

6 mg of protein was aliquoted for each sample. Sample volume was brought up to 20 mL in a solution of 4M Urea+50mM HEPES, pH=8.5. Disulfide bonds were reduced in 5mM dithiothreitol (DTT) for 30 minutes at 56°C. Reduced disulfide bonds were alkylated in 15mM of iodoacetamide (IAA) in a darkened room temperature environment for 20 minutes. The alkylation reaction was quenched via the addition of the original volume of DTT for 15 minutes in a darkened environment at room temperature. Proteins were next precipitated from solution via the addition of 5 µL of 100% w/v trichloroacetic acid (TCA). Samples were mixed and incubated on ice for 10 minutes. Samples were subjected to centrifugation at 16,000 x g for 5 minutes at 4°C. The supernatant was removed and sample pellets were gently washed in 50 µL of ice-cold acetone. Following the wash step, samples were subjected to centrifugation at 16,000 x g at 4°C. The acetone wash was repeated, and the final supernatant was removed. Protein pellets were dried on a heating block at 56°C for 15 minutes, and pellets were resuspended in a solution of 1M Urea+50mM HEPES, pH=8.5. The UPS2 standard (Sigma) was reconstituted as follows. 20 mL of a solution of 4M Urea+50mM HEPES, pH=8.5 was added to the tube. The sample tube was subjected to vortexing and water bath sonication for 5 minutes each. The standard was subjected to reduction and alkylation using methods described above. The sample was next diluted in a solution of 50mM HEPES, pH=8.5 such that the final concentration of Urea was 1M. 0.88 mg of the protein standard was spiked into each experimental sample, and samples were subjected to a two-step digestion process. First, samples were digested using 6.6µg of LysC at room temperature overnight, shaking. Next, protein was digested in 1.65 µg of sequencing-grade trypsin (Promega) for 6 hours at 37°C. Digestion reactions were terminated via the addition of 3.3 µL of 10% trifluoroacetic acid (TFA), and were brought up to a sample volume of 300 µL of 0.1% TFA. Samples were subjected to centrifugation at 16,000 x g for 5 minutes and desalted with in-house-packed desalting columns using methods adapted from previously-published studies^38,39^. Following desalting, samples were lyophilized, and then stored at −80°C until further use.

### LC-MS/MS

Samples were resuspended in a solution of 5% acetonitrile (ACN) and 5% formic acid (FA). Samples were subjected to vigorous vortexing and water bath sonication. Samples were analyzed on an Orbitrap Fusion Mass Spectrometer with in-line Easy NanoLC (Thermo) in technical triplicate. Samples were run on an increasing gradient from 6%-25% ACN+0.125% FA for 70 minutes, then 100% ACN+0.125% FA for 10 minutes. 1µg of each sample was loaded onto an in-house-pulled and -packed glass capillary column heated to 60°C. The column measured 30 cm in length, with outer diameter of 360 mm and inner diameter of 100 mm. The tip was packed with C4 resin with diameter of 5 mm to 0.5 cm, then with C18 resin with diameter of 3 mm an additional 0.5 cm. The remainder of the column up to 30 cm was packed with C18 resin with diameter of 1.8 mm. Electrospray ionization was achieved via the application of 2000 V to a T-junction connecting sample, waste, and column capillary termini. The mass spectrometer was run in positive polarity mode. MS1 scans were performed in the Orbitrap, with a scan range of 375-1500 m/z with resolution of 120,000. Automatic gain control (AGC) was set to 5 × 10^5^, with maximum ion inject time of 100 ms. Dynamic exclusion was performed at 30 second duration. Top n was used for fragment ion isolation, with n=10. The decision tree option was used for fragment ion analysis. Ions with charge state of 2 were isolated between 375-1500 m/z, and ions with charge state 3-6 were isolated between 600-1500 m/z. Precursor ions were fragmented using fixed Collision-Induced Dissociation (CID). Fragment ion detection occurred in the linear ion trap, and data were collected in profile mode. Target AGC was set to 1×10^4^. Technical triplicate spectral data was searched against a customized reference proteome comprised by the reference proteome of the respective strain (see above) appended to the UPS2 fasta sequences (Sigma) using Proteome Discoverer 2.1 (Thermo). Spectral matching and *in silico* decoy database construction was performed using the SEQUEST algorithm^40^. Precursor ion mass tolerance was set to 50 ppm. Fragment ion mass tolerance was set to 0.6 Da. Trypsin was specified as the digesting enzyme, and two missed cleavages were allowed. Peptide length tolerated was set to 6-144 amino acids. Dynamic modification included oxidation of methionine (+15.995 Da), and static modification included carbamidomethylation of cysteine residues (+57.021 Da). A false-discovery rate of 1% was applied during spectral searches.

### Protein abundance estimation

In order to estimate absolute protein abundance, the top3 metric was calculated for each protein as the average of the three highest peptide areas^11,12^. Robust linear regression (as implemented in the MASS package^41^) was used to calibrate top3 with the UPS2 standard according to the following model to obtain the amount of loaded protein A:

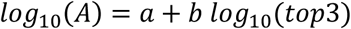

In order to obtain abundance relative to cell dry weight (C), we use a constant ratio γ = 13.94 µmol gDW^-1 42^:

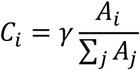

### Calculation of *k*_app,max_

For each biological replicate, apparent catalytic rates *k*_app_ were calculated as the ratio of protein abundance and measured flux if (1) the protein abundance surpassed 50 pmol gDW^-1^ and (2) the estimated flux was at least four times larger than the range of the 95% confidence interval, flux was larger than 100 fmol gDW-1 h-1, and the 95% confidence interval did not include zero, as defined in McCloskey et al.^29^.

For each of the two biological replicates per strain, *k*_app,max_ was calculated as the maximum *k*_app,max_ across the 21 strains. Finally the average *k*_app,max_ over the two replicates was calculated and used in the presented analyses.

### Parametric bootstrap for *k*_app,max_

We used a parametric bootstrap approach to estimate how experimental variability in proteomics and fluxomics data affects *k*_app,max_. We assumed protein abundance to be normally distributed with mean and standard deviation estimated from biological replicates. Variability in flux data was also assumed to take a normal distribution, where we used the standard deviation of the MFA procedure that resulted from multiple model reruns on biological triplicates^6–9^. For each reaction, 500 bootstrap samples were simulated, and these samples were used to calculate 95% confidence intervals for *k*_app,max_.

### Machine learning

Turnover numbers were extrapolated to the genome scale using the machine learning approach published previously^16^. The enzyme features used in Heckmann et al.^16^ were labelled with the *k*_app,max_ values estimated in this study, and an ensemble model of elastic net, random forest, and neural network was trained using the caret package^43^ and h2o^44^. Model hyperparameters were chosen in five times repeated cross-validation (one repetition in the case of neural networks) based on the RMSE metric, as reported in Heckmann et al.^16^. For the neural networks, random discrete search was used for optimization of hyperparameters^16^.

### MOMENT modeling

Validation of different turnover number vectors in the MOMENT model was conducted as described in Heckmann et al.^16^. The genome-scale metabolic model *i*ML1515^45^ was used in the R^46^ packages sybil^47^ and sybilccFBA^48^ to construct linear programming problems that were solved in IBM CPLEX version 12.7.

### ME modeling

To complement the MOMENT-based validation of the computed turnover numbers, a similar validation approach was employed with the *i*JL1678b-ME genome-scale model of *E. coli* metabolism and gene expression^49^. The *k*_app_s were mapped to *i*JL1678b-ME as previously described^16^. However the ME-model *k*_app_s were adjusted due to a key difference that lies in the way that the MOMENT and ME-model resource allocation models apply enzyme constraints. MOMENT accounts for each unique protein contained within a catalytic enzyme whereas the ME-model formulation accounts for the complete number of protein subunits in an enzyme. As a result, the macromolecular “cost” of catalyzing a reaction in the ME-model is often notably higher than in MOMENT. To account for this, the *k*_app_s in the ME-model were adjusted by scaling each *k*_app_ by the number of protein subunits divided by the number of unique proteins.

The ME-model was solved in quad precision using the qMINOS solver^50^ and a bisection algorithm^51^ to determine the maximum feasible model growth rate, within a tolerance of of 10^-12^. All proteins in a solution with a computed synthesis greater than zero copies per cell were compared to experimentally measured protein abundances. Since the ME-model accounts for the activity of many proteins outside of the scope of the *k*_app_ prediction method, only those that overlap with predicted *k*_app_s were considered.

## Supporting information

Supplementary Data 1: Table of mutations that occurred during ALE (csv)

## Data Availability

Results of genome sequencing and mutation calling were deposited to ALEdb (aledb.org) as part of the “Central Carbon Knockout (CCK) project”. Mass spectrometry-based proteomic data can be found on the ProteomeXchange Consortium (http://proteomecentral.proteomexchange.org) with the dataset identifier “PXD015344”.

## Acknowledgements

This work was supported by the Novo Nordisk Foundation Grant Number NNF10CC1016517. The authors would like to thank Dan Davidi and Douglas McCloskey for insightful discussions, and Marc Abrams for proofreading the manuscript.

## Author contributions

DH and BOP designed the study. DH, PP, and AMF conducted strain genotyping and strain selection. YH, AC, and MCT prepared samples for proteomics experiments. AC, DH, and DJG designed proteomics experiments. AC conducted LC-MS/MS experiments. CJL conducted ME modeling. DH conducted data analysis, data integration, training of machine learning models, and MOMENT modeling.

## Competing interests

The authors declare no competing interests.

## Additional Information

Supplementary Information 1: Supplementary Table 1 and Supplementary Figures (pdf).

Supplementary Data 1: Table of mutations that occurred during ALE (csv).

