## Supplementary Information 1: Supplementary Table 1 and Supplementary Figures (pdf) for "Kinetic profiling of metabolic specialists demonstrates stability and consistency of in vivo enzyme turnover numbers"

Supplementary Table 1: Correlation between biological replicates and coverage of proteomics samples. The $R^2$ of protein abundance on log scale between two biological replicates is shown along with the number of unique proteins that were detected in at least one of the two replicates.

| strain | $R^2$ between biological replicates | number of proteins detected |
|---|---|---|
| WT 1 | 0.92275 | 2105 |
| WT 2 | 0.818324 | 2066 |
| pgi 1 | 0.862347 | 2076 |
| pgi 2 | 0.928284 | 2158 |
| pgi 3 | 0.85155 | 2161 |
| pgi 4 | 0.904126 | 2117 |
| pgi 5 | 0.912196 | 2164 |
| pgi 6 | 0.919021 | 2017 |
| pgi 7 | 0.883619 | 2033 |
| pgi 8 | 0.884893 | 2045 |
| ptsHIcrr 1 | 0.920072 | 2132 |
| ptsHIcrr 2 | 0.908835 | 2138 |
| ptsHIcrr 3 | 0.92805 | 2129 |
| ptsHIcrr 4 | 0.897441 | 2159 |
| sdhCB 1 | 0.923292 | 1934 |
| sdhCB 2 | 0.920167 | 2033 |
| sdhCB 3 | 0.849117 | 1981 |
| tpiA 1 | 0.921315 | 2147 |
| tpiA 2 | 0.923444 | 2003 |
| tpiA 3 | 0.913019 | 1818 |
| tpiA 4 | 0.927285 | 1991 |

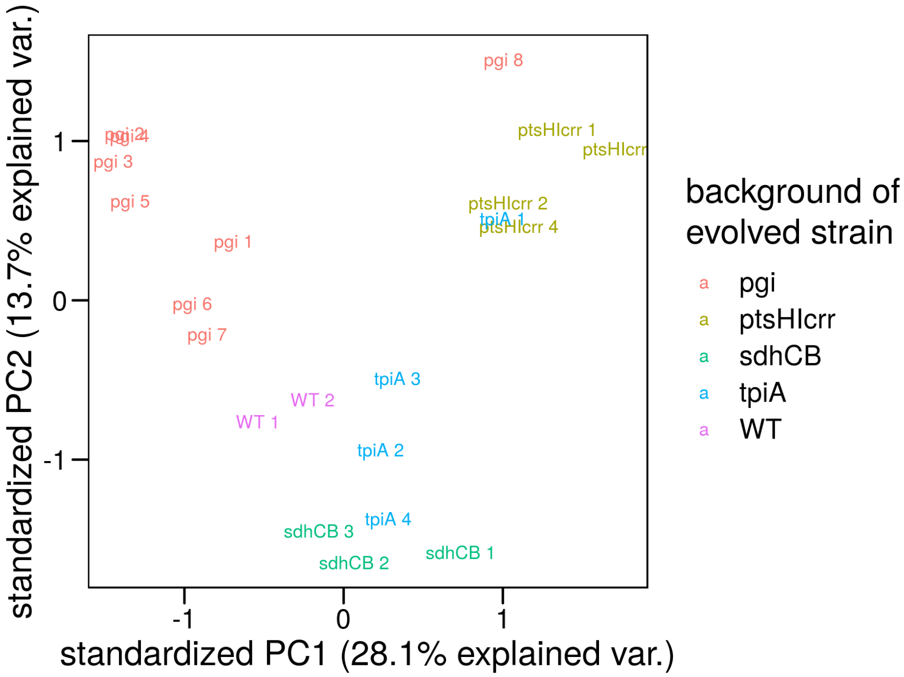


Supplementary Figure 1: PCA biplot of protein abundances. Protein abundances were log-transformed, centered, and scaled. Only proteins that were detected in all samples were used for this analysis (n = 829).

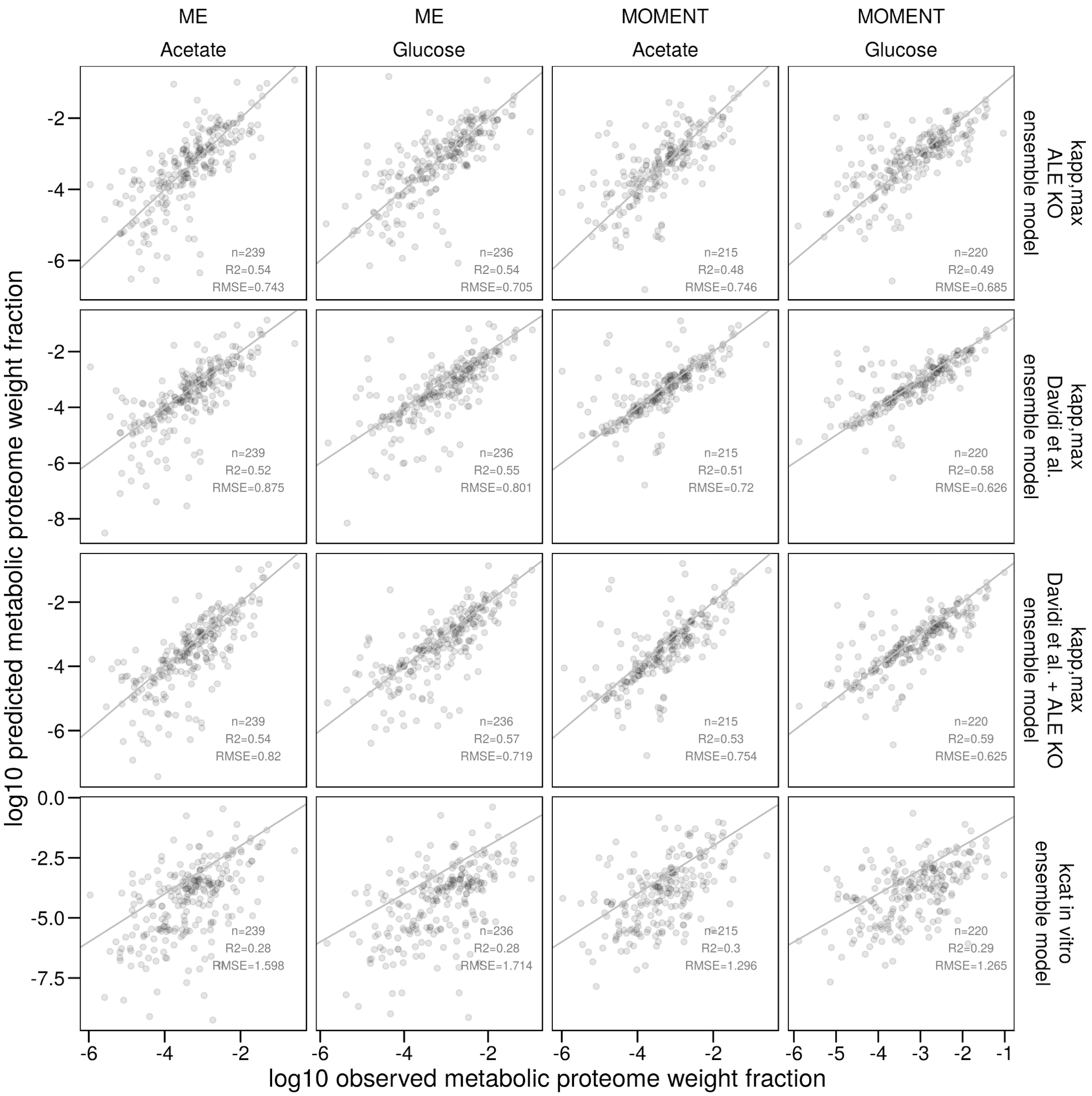


Supplementary Figure 2: Direct comparison of protein abundance predictions with measured data for different $k_{cat}$ parameterizations. Proteomics data for growth on glucoses and acetate from Schmidt et al.[1] is shown as examples
